# Probing site-specific RNA dynamics by solid-state NMR spectroscopy

**DOI:** 10.1101/2025.07.02.662307

**Authors:** Arun Kumar Sreemantula, John Kirkpatrick, Alexander Marchanka

**Affiliations:** Institute for Organic Chemistry and Centre of Biomolecular Drug Research (BMWZ), Leibniz University Hannover, Schneiderberg 38, 30167 Hannover, Germany; Laboratory of Integrative Structural Biology, School of Biosciences, College of Life and Environmental Sciences, University of Birmingham, Birmingham B15 2TT, UK; Department of Structure and Function of Proteins, Helmholtz Centre for Infection Research, Inhoffenstr. 7, 38124 Braunschweig, Germany; Goethe University Frankfurt, Institute for Biophysical Chemistry, Max von Laue Str. 9, 60438 Frankfurt am Main, Germany; Molecular Systems Biology Unit, European Molecular Biology Laboratory, Meyerhofstr. 1, 69117 Heidelberg, Germany

**Keywords:** solid state NMR, RNA-protein complex, dynamics, relaxation, MAS

## Abstract

Knowledge of site-specific dynamics of biomolecules is necessary to understand their specific functions. Solid-state NMR is a uniquely powerful technique that can provide both structural and motional information about complex biomolecules. In recent years, nucleic acids have become increasingly studied by ssNMR, but as yet few RNA or DNA structures are available, with ssNMR-derived dynamics data on RNA rarer still. Here, we report the first systematic ssNMR study of ^15^N *T*_1_ relaxation in RNA using straightforward nucleotide-type-specific and uniform labeling schemes. We observe clear correlation of the measured ^15^N *T*_1_ relaxation time-constants with different structural elements in the RNA molecule, reflecting the distinct characteristics of their underlying motions. We anticipate that this novel approach can be further developed to provide a detailed and comprehensive picture of RNA dynamics in large biomolecular machines.

## Introduction

Biomolecules undergo significant conformational changes in order to perform their various physiological functions.^1-3^ Characterizing this dynamic behavior is crucial for a proper understanding of the structure–function relationships that govern fundamental cellular processes such as allostery, protein-folding, signaling and enzymatic turnover. For many years, studies of biomolecular dynamics focused on proteins,^4-5^ as it was long thought that the biological role of RNA was restricted to that of a simple intermediary in the conversion of information from DNA to proteins. More recently, the outlook has changed as more of the myriad different roles of RNA have been discovered.^6-10^ It is now accepted that an in-depth understanding of RNA structure and function is critically important, particularly in the context of engineering

RNA molecules for use as drugs, vaccines,^11^ and tools for molecular and synthetic biology.^12-13^ The conformational flexibility of RNA makes its structural characterization by X-ray crystallography^14-15^ or cryo-electron microscopy^16^ challenging. In contrast, nuclear magnetic resonance (NMR) spectroscopy is ideally suited to investigations of flexible molecules, and has become the method-of-choice to study RNA structure and dynamics. Both of the two main types of NMR — solution and solid-state NMR — can be fruitfully applied to studies of RNA, but the applicability of solution NMR to RNAs of more than 100 nucleotides (nt) is hampered by the severe line-broadening associated with the slow rotational diffusion of large molecules in solution. Solid-state NMR (ssNMR) does not suffer from this problem, with magic-angle spinning (MAS) used to average anisotropic magnetic interactions, and can in principle be applied to complexes of any size by utilizing carefully chosen isotope-labeling schemes and recoupling experiments.^17-18^ To date, several ssNMR studies have addressed structural and dynamic features of membrane proteins,^19-20^ large molecular assemblies^21-22^ and amyloid fibrils.^23-24^ Despite these significant advances, investigations of RNAs and its complexes by ssNMR remain relatively few in number — the close chemical similarity of the four different nucleotides that constitute RNA and the correspondingly limited chemical-shift dispersion of the NMR signals represents a major challenge — and experimental methods for ssNMR of RNA are still in their infancy.^25-26^ Nonetheless, a few research groups, including our own, have made significant recent progress in the study of structure and function of RNA and RNP complexes by ssNMR, exemplified by the first structures of an RNA and a protein–RNA complex,^27-28^ studies of dimerization in HIV-1 RNA,^29^ characterization of an adenine riboswitch^30^ and characterization of a very large native RNA from the bacteriophage MS2.^31^ Furthermore, utilization of the sensitivity enhancement by DNP allowed to obtain insights into structures of protein-RNA,^32^ RNA-RNA-protein complexes^33^ and nucleic acids in native environment.^34^ However, ssNMR studies of RNA dynamics have so far been limited mostly to the efforts of the Drobny group, whose work includes characterizing motions of HIV-TAR RNA using deuterium relaxation and analysis of deuterium line-shapes on specifically 5,6-^2^H-labeled pyrimidine nucleotides.^35-37^ Very recently, Wang group has applied water-RNA exchange spectroscopy in ssNMR to obtain insights into stability of base-pairs.^38^ Despite very little number of studies, ssNMR spectroscopy holds great potential for studies of RNA dynamics, partly due to the complete absence of the averaging mediated by rotational diffusion in solution, which should allow the characterization of dynamics over a wide and uninterrupted range of timescales from picoseconds to seconds, including the nanosecond–microsecond “blind-spot” that is mostly inaccessible by solution NMR techniques.^39^

Here, this work presents an innovative and simple yet effective ssNMR approach for quantifying the dynamics of nucleotide-type-specifically and uniformly labeled RNA. We report for the first time ssNMR-derived ^15^N longitudinal relaxation data from RNA — acquired on micro-crystalline 26mer box C/D RNA in complex with the protein L7Ae^27^ — revealing both nucleotide-specific and atom-specific relaxation time-constants (*T*_1_s). Relaxation data acquired at different MAS rates provides insights into the influence of spin-diffusion on the measurement of ^15^N *T*_1_ relaxation time-constants in RNA and builds towards a standardized framework for characterization of RNA dynamics by ssNMR relaxation measurements.

## Materials and Methods

### Sample preparation

The complete procedure for preparation of L7Ae–Box C/D 26mer RNA complex for solid-state NMR studies has been described in detail elsewhere.^28, 40^

In total, four different RNP samples were prepared for this study:

- Three nucleotide-type-specifically labeled samples for measurements at 16 kHz and 55 kHz MAS at 600 MHz magnetic field:
  1. A^lab^ RNA: 3 mg ^13^C,^15^N A^lab^ 26mer box C/D RNA + 5 mg unlabeled L7Ae protein.
  2. G^lab^ RNA: 2.5 mg ^13^C,^15^N G^lab^ 26mer box C/D RNA + 3.7 mg unlabeled L7Ae protein.
  3. U^lab^ RNA: 1.9 mg ^13^C,^15^N U^lab^ 26mer box C/D RNA + 2.8 mg unlabeled L7Ae protein. These three samples were first packed into 3.2-mm standard-wall rotors for 16 kHz MAS experiments and then repacked into 1.3-mm rotors for 55 kHz MAS experiments.
- 300 μg uniformly ^13^C,^15^N-labeled 26mer box C/D RNA + 500 μg unlabeled L7Ae protein.

This sample was packed into a 0.81-mm rotor for 100 kHz and 55 kHz MAS experiments at a 850 MHz magnetic field. The same sample was used for the ^1^H resonance assignments and base-pair-pattern elucidation reported in our previous work.^40^

### NMR spectroscopy

A Bruker Avance III HD spectrometer operating at a ^1^H Larmor frequency of 600 MHz and equipped with standard 3.2-mm and 1.3-mm triple-resonance ^1^H/^13^C/^15^N MAS probeheads (Bruker Biospin) was utilized to perform NMR experiments at 16 kHz and 55 kHz MAS, respectively.

A Bruker Avance III HD spectrometer operating at a ^1^H Larmor frequency of 850 MHz and equipped with a 0.81-mm triple-resonance ^1^H/^13^C/^15^N MAS probe head developed in the Samoson laboratory (https://www.nmri.eu/)^41^ was utilized to perform NMR experiments at 100 kHz and 55 kHz MAS.

For the collection of *T*_1_ relaxation time constants at 16 kHz MAS ^13^C detected experiments were utilized, while for the data collection at 55 kHz and 100 kHz MAS ^1^H detected experiments were utilized.

After initial proton *π*/2 pulse, ^1^H polarization is transferred to directly attached ^15^N using short CP step. Then the transverse ^15^N magnetization is rotated to the longitudinal axis and allowed to relax during a variable delay *τ*, before being returned to the transverse plane for an indirect chemical-shift evolution period. Finally, the ^15^N magnetization is transferred by CP to either ^13^C (^13^C-detected experiments, Figure S1A) or ^1^H (^1^H-detected experiments, Figure S1B) for detection. Typical ^1^H and ^15^N 90° pulse lengths for spectra acquired using 3.2 mm probehead were 2.8 μs and 6.0 μs, respectively. Typical ^1^H and ^15^N 90° pulse lengths for spectra acquired using 1.3 mm probehead were 1.5 μs and 3.8 μs, respectively, while those for spectra acquired using 0.81 mm probehead were 1.3 μs and 3.0 μs, respectively. All spectral acquisition parameters and details of each CP step are provided in tables S1 and S2. NMR pulse sequences are provided in Figure S1.

### Data collection, processing & analysis

Most of the 2D planes corresponding to different relaxation delays were acquired in an interleaved fashion, with a subset of pre-defined *τ* delays (table S1), resulting in pseudo-3D ^13^C,^15^N or ^1^H,^15^N spectra. Multiple pseudo-3D spectra (labeled below with the index *j*) corresponding to different subsets of relaxation delays were acquired separately due to the restrictions imposed by field-drift and cryogen refills. The full collection of 2D planes extracted from the pseudo-3D spectra and corresponding to the entire set of relaxation delays represents a complete relaxation data-set from which intensities were extracted for regression analysis.

^13^C chemical-shifts were referenced as described by Morcombe and Zilm.^42 15^N and ^1^H chemical-shifts were referenced indirectly using the chemical-shift referencing ratios from the work of Markley *et al*.^43^

#### Spectra processing and peak-fitting

Spectra were processed in Topspin using parameters given in Table S3 and analyzed with CcpNmr Analysis (v2.4).^44^ The assignments from our recent work^27, 40^ were used to assign the peaks, and peak-intensities were extracted as box-sum integrals using the box-sizes reported in table S3. *T*_1_ relaxation time-constants were determined using a global fitting procedure in OriginPro 8 software,^45^ whereby the peak-intensities from multiple pseudo-3D datasets (without any prior normalization between data-sets) were fitted to the mono-exponential decay function (1):

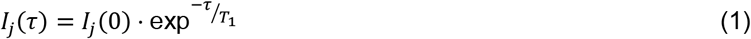

*I*_*j*_(*τ*) is the peak-intensity at delay *τ* from pseudo-3D spectrum *j, I*_*j*_ (0) is the peak-intensity at zero delay in spectrum *j* and *T*_1_ is the relaxation time-constant (in seconds). Normalized experimental data-points *I*_*j*_(*τ*)/*I*_*j*_(0) were plotted together with the mono-exponential decay fit in OriginPro 8.^45^

## Results and Discussion

The local flexibility of the polypeptide backbone of proteins can be accurately probed with solution NMR by measuring the ^15^N *T*_1_ and *T*_2_ time-constants and the {^1^H}^15^N steady-state NOEs of the individual amide groups,^46-47^ and with ssNMR first insights into flexibility can be obtained by measuring the ^15^N *T*_1_s alone.^48-51^ In contrast, for RNA there are no protonated nitrogens of the same chemical type that are present in all four nucleotides. This complicates systematic characterization of RNA dynamics from solution-state ^15^N relaxation data, which is only straightforward for base-paired imino nitrogens.^52^ ssNMR can detect both base-paired and non-base-paired nitrogens, and is also more suited to direct observation of non-protonated nitrogen atoms N1 and N9 in pyrimidines and purines, respectively. However, measurement of relaxation data from these nitrogens is impractical due to the long ^1^H-^15^N cross-polarization (CP) periods^53-54^ that are required (> 2 ms), which lead to weak signals and hence unacceptably long experimental times. This restricts the measurable ^15^N *T*_1_ relaxation time-constants to those of protonated nitrogens in imino (NH) and amino (NH_2_) groups. For these nuclei, efficient generation of ^15^N magnetization can be accomplished with a short ^1^H-^15^N CP step (<1 ms) (Figure S1A–B). Importantly, the two types of nitrogen (imino and amino) are differently distributed amongst the four nucleotides, with imino groups present in guanosine (N1–H1) and uridine (N3–H3), and amino groups in adenosine (N6–H61/H62), cytidine (N4–H41/H42) and guanosine (N2–H21/H22) (Figure 1A). For similar dynamics, the *T*^1^ relaxation time-constants of amino nitrogens are expected to be shorter than those of imino nitrogens due to the two one-bond N–H dipolar interactions in the former compared to the single interaction in the latter.^55^ This means that amino (A-N6, C-N4, G-N2) and imino (G-N1, U-N3) relaxation data must be considered separately to build an accurate picture of the sequence-specific dynamics within an RNA molecule. In addition, the intrinsic relaxation characteristics of the imino nitrogens of guanosine and uridine are likely to differ slightly due to the dipolar interactions between G-N1 and the protons of G-N2 (G-H21/H22); interactions that are not present in uridine.

**Figure 1.**
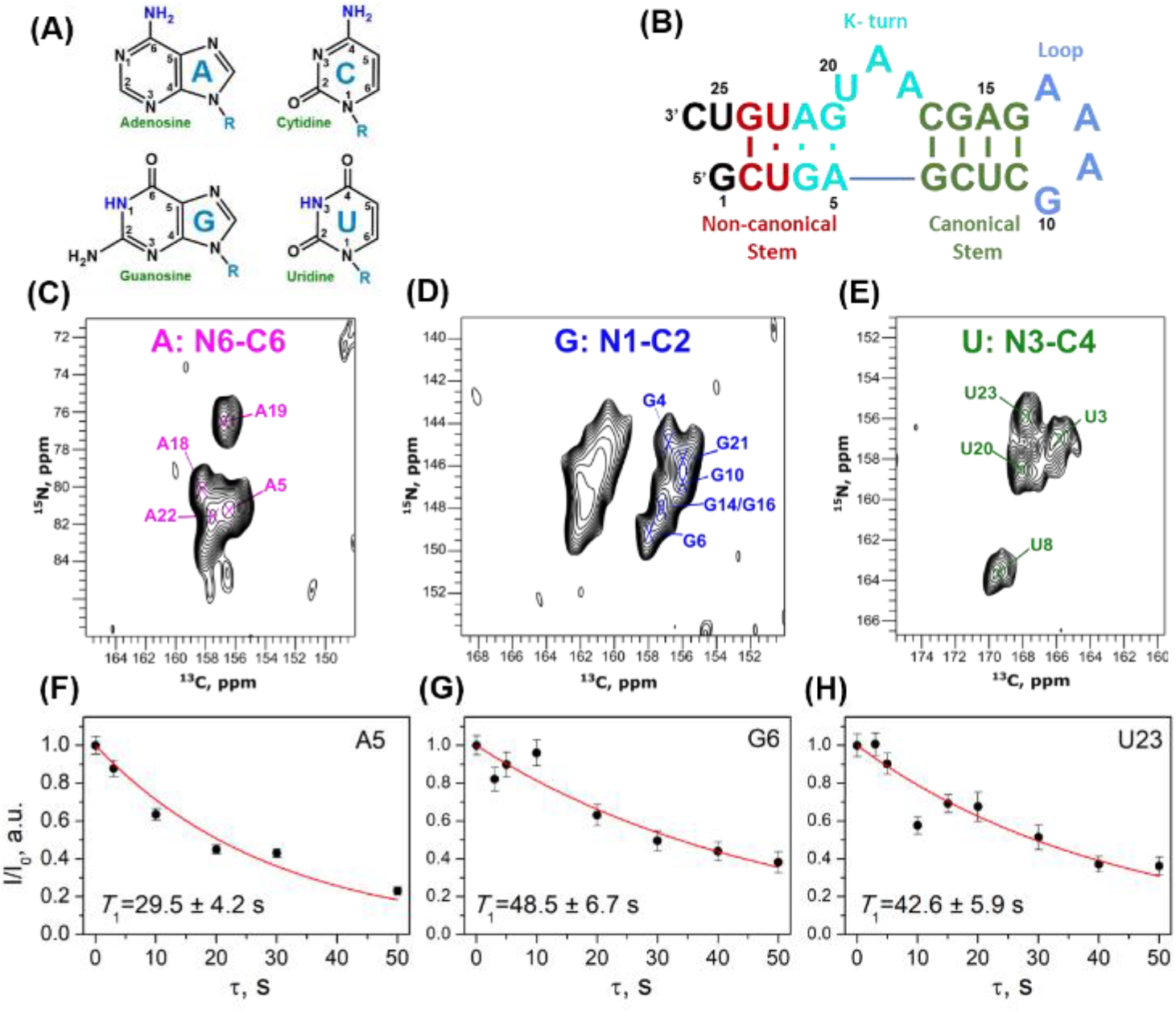
Relaxation data acquired at 16 kHz MAS. **(A)** Nucleobase structure of four nucleotides typically present in RNA. **(B)** Secondary structure of 26mer box C/D RNA used in this study. **(C–E)** 2D NC correlation spectra corresponding to zero relaxation delay for A^lab^-RNA **(C)**, G^lab^-RNA **(D)** and U^lab^-RNA. **(E)**. The unassigned peak-group in (D) belongs to overlapped N1-C6 correlations. **(F–H)** Examples of relaxation decay curves for nucleotides A5 **(F)**, G6 **(G)** and U23 **(H)**. Experimental data-points are in black, and mono-exponential decay fits are shown in red. All data was acquired on nucleotide-type-specifically labeled 26mer box C/D RNA in complex with L7Ae protein at a field-strength of 600 MHz.

Due to absence of any previous work on measuring ^15^N *T*_1_ relaxation of RNA by ssNMR, and to establish a firm foundation for a standardized framework of such relaxation studies, we decided to acquire *T*_1_ relaxation data over a range of MAS rates, utilizing 3 different probeheads (3.2 mm, 1.3 mm and 0.8 mm). While with 3.2 mm rotors only heteronuclear (^13^C) detection experiments were feasible, 1.3 mm and 0.8 mm rotors have allowed high-sensitive ^1^H detection. *Vice versa*, low sensitivity heteronuclear detection with 1.3 mm and 0.8mm rotors was not feasible due to low amount of packed sample and therefore, low signal intensity. We have built our work on the pioneering approach of the Emsley group in their ^15^N *T*_1_ relaxation study of the crystalline protein Crh by ssNMR at 10 kHz MAS,^48^ and have also drawn on the work of Giraud *et al*. in estimating the influence of spin-diffusion on the observed ^15^N longitudinal decay curves.^56^ In brief, we measured the site-specific ^15^N longitudinal relaxation time-constants (*T*_1_s) of 26mer box C/D RNA in complex with L7Ae protein (Fig. 1B) at multiple (16, 55, 100 kHz) MAS rates using either uniformly or nucleotide-type-specifically ^13^C,^15^N-labeled RNA samples. These specific MAS rates were chosen as safe rotation speeds for 3.2 mm, 1.3 mm and 0.81 mm rotors, respectively.

### ^15^N-*T*_1_ relaxation in RNA at 16 kHz MAS (600 MHz field-strength)

To achieve sufficient spectral resolution at low-to-moderate MAS rates (3.2-mm rotor-diameter; 16 kHz MAS), ^15^N *T*_1_ relaxation time-constants were extracted from ^13^C-detected 2D ^15^N−^13^C correlation spectra on nucleotide-specifically ^13^C,^15^N-labeled (A^lab^, G^lab^, U^lab^) 26mer Box C/D RNA (Figure 1B). We did not attempt measurement of C^lab^ 26mer RNA due to the significant spectral overlap that we knew to exist with this labeling scheme.^57^ “Zero-delay” spectra (acquired with a relaxation delay of zero) are shown in Figure 1C– E, while magnetization-transfer schemes and pulse sequences are shown in Figure S1A. Resonance assignment and fitting of peak intensities to extract *T*_1_ values was achievable for 13 of the 26 nucleotides in the molecule. In total, *T*_1_ relaxation data was analyzed for amino nitrogen N6 in adenosines A5, A18, A19 and A22, imino nitrogen N1 in guanosines G4, G6, G10, G14/G16 and G21, and imino nitrogen N3 in uridines U3, U8, U20 and U23. While the peaks of the adenosines and uridines are well-resolved, those of guanosines are less so, with heavy overlap of G14 and G16 and partial overlap of G4, G10 and G21, so that only semi-quantitative analysis for these nucleotides was possible. Examples of the intensity decay curves and corresponding mono-exponential fits for nucleotides A5, G6 and U23 are shown in Figure 1F– H, while the intensity decay-profiles and fits for all analyzed nucleotides are shown in Figure S2. The extracted apparent relaxation time-constants vary over quite a broad range, from 12.8 s up to 50.5 s (Supplementary Table S5). A significantly longer average time-constant is observed for imino nitrogens than for amino nitrogens; the average imino *T*_1_ (guanosines and uridines, 9 datasets) was 38.5 s (with a standard deviation (s.d.) of 7.0 s), while the average amino *T*_1_ (adenosines, 4 datasets) was 22.1 s (s.d. 9.5 s). This difference in the average *T*_1_ for imino and amino nitrogens is explained by the difference in the associated spin-system architectures. In addition to these expected nucleotide-type-specific differences, site-specific variations are also observable. Amino nitrogens of adenosines located in base-paired regions (A5 and A21) have an average *T*_1_ value of 30.2 s, while those of two others located in the flexible kink-turn region (A18 and A19) have an average value of 14.0 s. However, no significant variations are observed between different guanosines and uridines, despite these two types of nucleotides being distributed across different structural regions of the RNA.

### ^15^N-*T*_1_ relaxation in RNA at 55 kHz MAS (600 MHz field-strength)

Next, we proceeded to acquire relaxation data at 55 kHz and 100 kHz MAS (1.3-mm and 0.81-mm rotor-diameters, respectively). At these MAS rates, ^1^H-detection on RNA becomes feasible,^58^ so that we were able to follow decay of longitudinal ^15^N magnetization in ^1^H,^15^N CP-HSQC-type spectra. The measurements at 55 kHz MAS were made on the nucleotide-type-specifically labeled samples previously employed for the 16 kHz measurements, while a single uniformly ^13^C,^15^N-labeled sample was used for the measurements at 100 kHz MAS. We have recently reported the resonance assignment of the imino- and amino-protons in this 26mer RNA.^40^ Panels A–C of Figure S3 show representative CP-HSQC spectra of the amino region of A^lab^ and the imino regions of G^lab^ and U^lab^ 26mer box C/D RNA in complex with L7Ae protein, while panel D shows the experimental relaxation decay curves and corresponding mono-exponential fits.

Similar to the picture at 16 kHz MAS, at 55 kHz MAS the amino nitrogens have a significantly shorter average *T*_1_ value (28.6 s with s.d. of 13.6 s) than the imino nitrogens (63.2 s with s.d. of 11.7 s). The *T*_1_ values of the amino nitrogens of A18 and A19 (17.5 ± 1.9 s and 20.7 ± 1.4 s) are noticeably shorter than those of A5 and A22 (28.2 ± 2.2 s and 47.8 ± 4.9 s), indicating significantly increased mobility for those amino-nitrogens in non-base-paired regions. The imino nitrogens of all uridines except U20 have *T*_1_ values ≥ 58 s, while the bulged-out U20 has a *T*_1_ value of 45.1 ± 2.5 s. This funding for U20 is surprising, since while not base-paired, U20 still should have limited flexibility due to the direct contact of U20 with the cognate L7Ae protein. The imino nitrogens of base-paired guanosines (G4, G6, G14/G16, G21) have *T*_1_ values that are on average 60% longer than that of G10 (located in the loop region).

For various reasons, the relaxation data for 13 of the 26 nucleotides of the box C/D RNA could not be reliably measured at 16 kHz and 55 kHz MAS. While peaks from nucleotides G1, A11, A12, A13, U25 and C26 were never observed in any of our experiments due to the high flexibility of the terminal regions and broad peaks of adenosines residues in the GAAA tetraloop likely resulting from a partial lack of defined orientation/alignment during sample preparation, leading to local disorder, the resonances of nucleotides C2, C7, C9, A15, C17 and G24 have been assigned in our previous studies.^27, 40, 59^ Unfortunately, they suffer from low signal-intensity and their peaks tend to be heavily overlapped, so that extraction of reliable relaxation data at 16 kHz and 55 kHz MAS was not possible. However, we were able to acquire relaxation data for the cytidines of this group at 100 kHz MAS, due to the significantly improved spectral resolution and sensitivity of these experiments (*vide infra*).

The *T*_1_ values of both amino and imino nitrogens at 55 kHz MAS are significantly longer than the corresponding values acquired at 16 kHz MAS. The experiments at 16 kHz and 55 kHz MAS were both conducted at 600 MHz, so that the *true T*_1_ values — which depend only on the strength of the dipolar interactions and the spectral densities at the frequencies of the nuclear-spin transitions — should be the same. That shorter *T*_1_ values are extracted at 16 kHz MAS is almost certainly the result of non-negligible spin-diffusion at the slower MAS rate, which will impact the measured *apparent* relaxation time-constants.^39^ While a study of ^15^N longitudinal relaxation in proteins at 10 kHz MAS^56^ showed that the impact of spin-diffusion on the apparent relaxation time-constants should be minimal for amide nitrogens in a polypeptide chain, the effect for nitrogen nuclei in RNA was never studied. In proteins, amide nitrogens in two adjacent residues are three bonds apart, with typical N–N distances of ∼3 Å, while most nitrogen-pairs in RNA nucleobases have only two intervening bonds and hence are more closely separated. We acquired 2D ^15^N,^15^N proton-driven spin-diffusion^60^ (PDSD) spectra at 16 kHz MAS with a mixing-time of 10 s, which revealed intense intranucleotide cross-peaks for guanosines and uridines, indicating the presence of significant N–N spin-diffusion in these nucleotides under slow MAS conditions (Figure S6B–C).^61^

We therefore made approximate estimations of the rates of N–N spin-diffusion in RNA nucleobases (see Supplementary Information for details) based on the approach introduced by the Emsley lab^56, 62^ in order to confirm that the effect of spin diffusion is responsible for the shortening of the apparent *T*_1_ values at 16 kHz MAS compared to those at 55 kHz MAS. Briefly, while the measured magnetization exchange rates in 26mer RNA in complex with L7Ae protein at 16 kHz MAS (typically 0.01–0.025 s^−1^) are similar to those measured in proteins at 10 kHz MAS, the auto-relaxation rates of the nitrogen nuclei in 26mer RNA are significantly slower than for those in proteins, which means that the distortion of the ^15^N magnetization decay curves due to spin-diffusion will be more pronounced for RNA. Simulations of the ranges for the apparent *T*_1_ values at 16 kHz MAS using *T*_1_ values at 55 kHz MAS and variable exchange rates as input parameters confirms that these apparent *T*_1_^app^ (16 kHz) values are always shorter than the true *T*_1_ values and are in good agreement with experimentally measured *T*_1_s at 16 kHz MAS (Table S5 and Figure S7). Therefore, the *T*_1_ values extracted from decay curves measured at 16 kHz MAS are erroneously shortened due to the effects of spin diffusion, with the most pronounced effect for the slowly relaxing imino nitrogens. Consequently, quantitative analysis of ^15^N *T*_1_ relaxation data acquired at 16 kHz MAS is highly challenging and would require accurate measurements of the various spin-diffusion-mediated exchange rates.

The reduction of spin-diffusion that is therefore required to facilitate acquisition of quantitative ^15^N *T*_1_ relaxation data can be achieved using a few different approaches. While uniform deuteration and site-specific labeling will dilute the dipolar-coupled spin-network and provide improved data-quality at lower MAS rates, these labeling approaches come at a cost of more expensive and tedious sample preparation. In contrast, performing the measurements at increased MAS rates yields pure relaxation decay curves, due to the near-complete removal of N-N spin-diffusion that results from more efficient averaging of anisotropic dipolar interactions, and can be used for nucleotide-type-specifically or even uniformly labeled samples.^63^ In summary, this investigation into the effects of spin-diffusion at low MAS rate of 16 kHz leads to the recommendation that quantitative ^15^N *T*_1_ relaxation data should always be collected at significantly high (≥ 55 kHz) MAS rates. Furthermore, dipolar cross-relaxation effects complicate the interpretation of ^15^N relaxation in the amino (NH_2_) nitrogens of adenosines, and this phenomenon can be quite complex. A detailed estimation and mechanistic understanding would require further in-depth analysis, which is beyond the scope of this paper.

### ^15^N-*T*_1_ relaxation in RNA at 100 kHz MAS (850 MHz field-strength)

Due to the superior spectral quality under ultra-fast MAS (100 kHz) and at high field-strength (850 MHz), a single sample of uniformly ^13^C,^15^N-labeled RNA was sufficient to acquire a full set of ^15^N *T*_1_ relaxation data. Previously, the same sample had been used to assign the nucleobases and determine the pattern of base-pairing in this RNA.^40^ The representative 2D ^1^H,^15^N CP-HSQC spectra of the amino and imino regions of the 26mer box C/D RNA in complex with L7Ae protein acquired under these conditions are shown in Figures 2A and 2B, respectively. While the imino region is well-resolved and allows straightforward extraction of the relaxation data, the amino protons of adenosines, cytidines and guanosines show significant line-broadening due to strong ^1^H,^1^H dipolar interactions, complicating extraction of signal intensities for relaxation data-analysis. However, compared to the spectra at 55 kHz MAS, the improved resolution at 100 kHz MAS means that the N4–H4 correlations in cytidines are resolvable and determination of their ^15^N-*T*_1_s becomes feasible. Overall, at 100 kHz MAS, relaxation data was analyzed for the amino nitrogens of adenosines and cytidines and for the imino nitrogens of guanosines and uridines.

**Figure 2.**
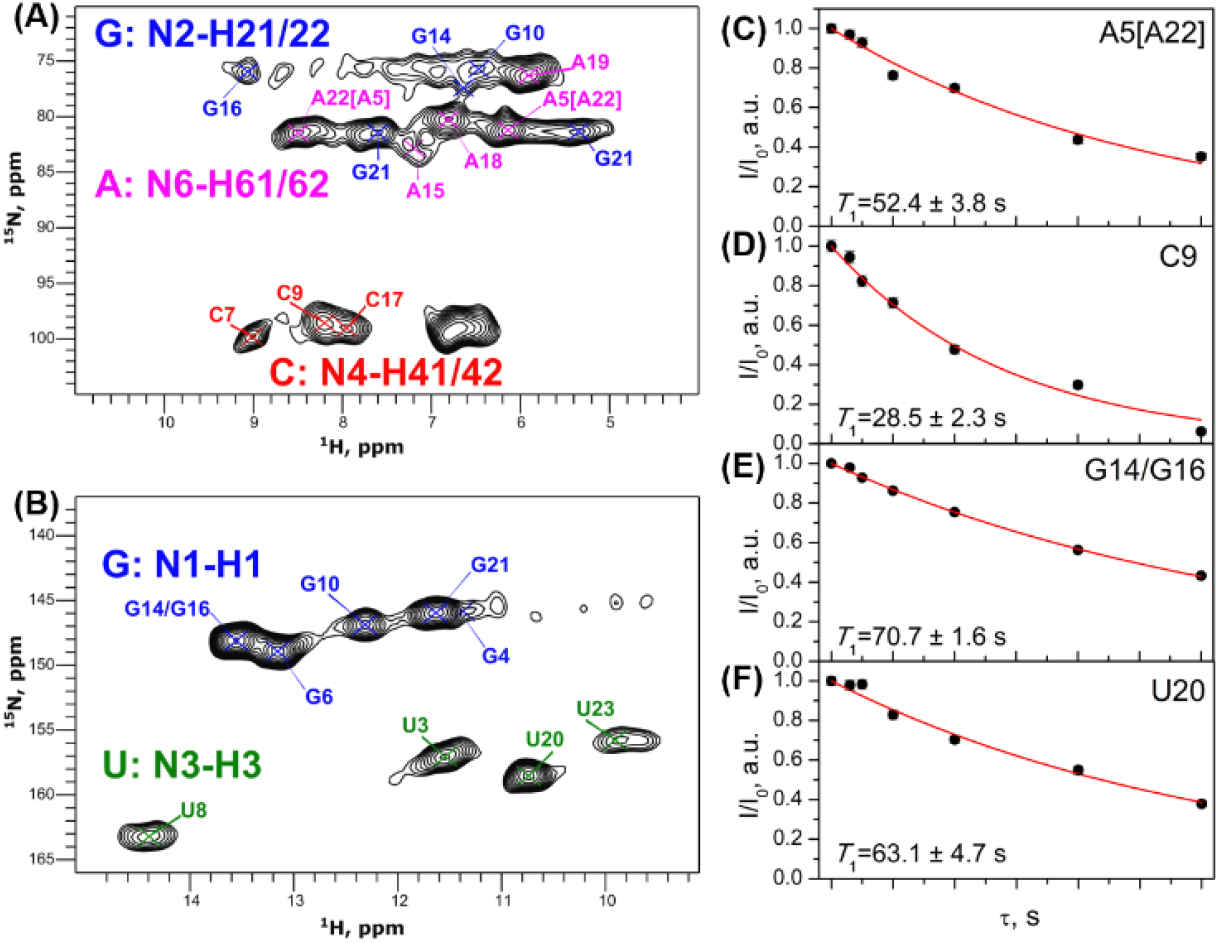
Relaxation data acquired at 100 kHz MAS. **(A–B)** 2D CP-HSQC-type ^1^H,^15^N correlation spectra acquired at zero relaxation delay showing amino **(A)** and imino **(B)** regions. Due to the near-degeneracy of the respective ^15^N chemical shifts, the relative assignment of the two amino peaks of A5 and A22 could not be determined with complete confidence; the labels A5[A22] and A22[A5] indicate the more-probable[less-probable] relative assignment. **(C–F)** Examples of relaxation decay curves for nucleotides A5[A22] **(C)**, C9 **(D)**, G14/G16 **(E)** and U20 **(F)**. Experimental data-points are in black and mono-exponential decay fits are in red. All data was acquired on uniformly ^13^C,^15^N labeled 26mer box C/D RNA in complex with L7Ae protein at a field-strength of 850 MHz.

The experimental decay curves show good fits to mono-exponential decay functions; examples for nucleotides A5/A22, C9, G14/G16 and U20 are shown in Figure 2C–F, while all the analyzed datasets are shown in Figure S4.

Due to the different numbers of directly bonded protons, and as previously observed at the lower MAS rates, the *T*_1_ time-constants for amino nitrogens (39.4 s with s.d 15.1 s) are significantly shorter than those for imino nitrogens (74.9 s with s.d. 17.5 s). The *T*_1_ time-constants of many nucleotides are significantly longer at 100 kHz MAS than at 55 kHz MAS. While we can assume negligible influence of spin-diffusion on ^15^N *T*_1_ values at MAS rates above 50 kHz (see supplementary Figure S7), the relaxation data at 100 kHz MAS was acquired at a higher magnetic field-strength than that at 55 kHz MAS (850 MHz *vs*. 600 MHz), which changes the absolute frequencies of the relevant nuclear-spin transitions.^64-65^ This means that the extracted relaxation rates are reporting on different parts of the spectral-density function that describes the dynamic behavior of the associated tensorial interactions. Since the spectral-density function is always a decaying function of frequency, it follows that relaxation time-constants that depend only on spectral-densities at non-zero frequencies will generally become longer when measured at higher magnetic field-strengths. Indeed, *T*_1_ relaxation time constants measured at 55 kHz Mas at 850 MHz magnetic field are in perfect agreement with 100 kHz MAS values (supplementary Figure S7B and Table S5).

As previously observed at slower MAS rates, the amino nitrogens of nucleotides A18 and A19 have significantly shorter relaxation time-constants (24.4 ± 3.6 s and 30.9 ± 2.9 s, respectively,) than nucleotides A5[A22] and A22[A5] (52.4 ± 3.8 s and 67.0 ± 9.4 s, respectively; see also legend to Figure 2A–B), which is due to increased flexibility in the k-turn (see Figure 3A). Despite all the cytidines being located in the canonical stem, the *T*_1_ of C9 is ∼30% shorter than the *T*_1_s of C7 and C17, which may be due to the proximity of C9 to the flexible tetraloop. Similarly, G10 — the first residue of the tetraloop — shows a significantly shorter *T*_1_ (50.1 ± 9.0 s) than those of nucleotides G6 and G21 (∼100 s), which are located in base-paired regions of the RNA. The peak assigned to the G14/G16 pair yields an intermediate *T*_1_ value of 70.7 ± 1.6 s, but due to the overlap this of course represents a pseudo-average of the individual *T*_1_s of G14 and G16 (Figure 2B). All uridine imino nitrogens have *T*_1_s ≥ 60 s. U3 and U23 in the non-canonical stem show slightly longer *T*_1_s (∼70 s) than those of nucleotides U8 and U20 (∼60 s). Interestingly, residue U20, which at 55 kHz MAS appears to relax more quickly than the other uridines, does not show such behavior at 100 kHz MAS.

**Figure 3.**
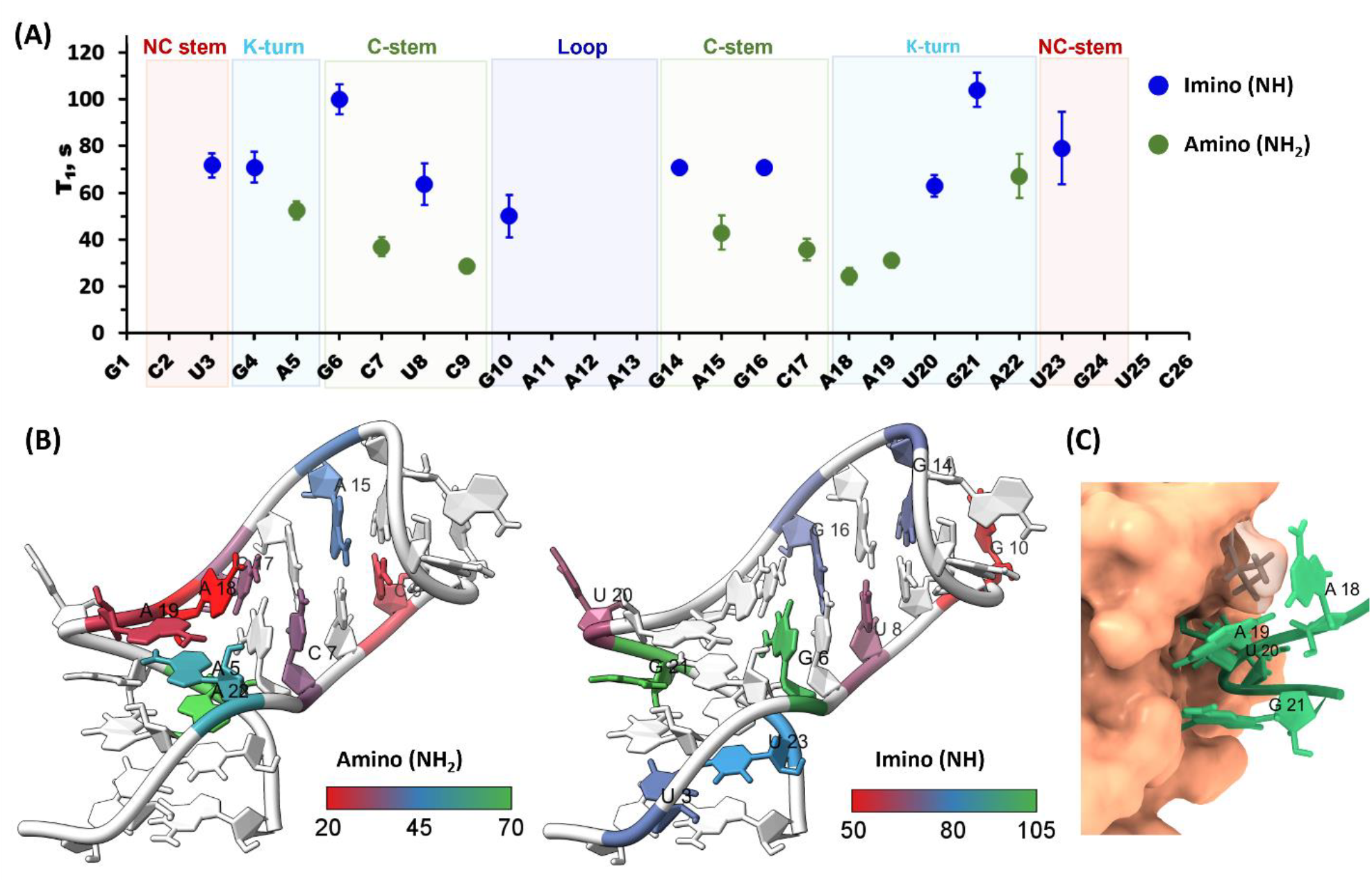
Site-specific ^15^N *T*_1_ relaxation of the 26mer box C/D RNA in complex with L7Ae protein. **(A)** *T*_1_ relaxation time-constants for amino (green) and imino (blue) nitrogens in uniformly ^13^C,^15^N labeled 26mer box C/D RNA measured at 100 kHz MAS and a field-strength of 850 MHz. The secondary-structure elements are indicated at the top. **(B)** Color-map representation of the amino (left) and imino (right) relaxation time-constants on the structure of the 26mer box C/D RNA^27^. **(C)** Close-up view of the protein–RNA interface showing the nucleotides involved in close protein–RNA contacts (the L7Ae protein is shown in surface representation).

Overall, the ^15^N *T*_1_ relaxation data obtained at 55 and 100 kHz MAS shows a clear sequence-specific pattern over the RNA molecule. Nucleotides located in non-base-paired regions of the RNA (C9, G10, A18 & A19 in the loop and k-turn) show faster-than-average ^15^N *T*_1_ relaxation (Fig. 3A–B), while base-paired nucleotides, both in the stems and the k-turn, are characterized by slower ^15^N relaxation (indicative of more limited flexibility in these regions). Base-paired nucleotides in the k-turn (G4:A22 and G21:A5) show the longest ^15^N *T*_1_ relaxation time-constants of all the nucleotides. Interestingly the non-base-paired and bulged-out U20 does not show the faster ^15^N *T*_1_ relaxation due to its stabilization in the prominent kink-turn due to the direct contact of U20 with the cognate L7Ae protein (Fig. 3C).

While ^15^N *T*_1_ relaxation in proteins has been extensively studied by ssNMR,^48-50, 62^ there have previously been no measurements of ^15^N *T*_1_ relaxation data for RNA in the solid-state. In this study, we show that ^15^N *T*_1_s in microcrystalline RNA are significantly longer than in proteins, while the residue-to-residue variation is noticeably smaller. The *T*_1_ values in the protein Crh differ by a factor of seven along the backbone, with shortest and longest time-constants of 7.1 s and 50 s, respectively, and with average value of 18.9 s.^48^ Similarly, for Gb1, the longest and shortest *T*_1_ values differ by a factor 40, with shortest and longest time-constants of 2.4 s and 91 s, respectively (average value of 23.8 s).^50^ The *T*_1_ variation (difference between shortest and longest *T*_1_s) in the 26mer box C/D RNA corresponds to a factor of 2.7 for amino nitrogens (A18 and A5/A22) and a factor of 2 for imino nitrogens (G10 and G21). Unfortunately, we were unable to measure relaxation data for residues A11–A13 in the tetra-loop, where some of these residues are expected to exhibit significantly increased mobility relative to the remainder of the molecule.

## Conclusion

In this study, we have presented for the first time ^15^N *T*_1_ relaxation data on RNA in complex with protein by MAS ssNMR, using easy-to-prepare nucleotide-type-specifically or uniformly labeled samples. We acquired relaxation data at three different MAS rates (16, 55 and 100 kHz, which are safe rotation rates of 3.2 mm, 1.3.mm and 0.8 mm MAS rotors, respectively). Relaxation data acquired at three different MAS rates revealed that spin-diffusion significantly perturbs the longitudinal ^15^N magnetization decay curves of protonated RNA at 16 kHz MAS, with the result that ^15^N *T*_1_ time-constants can be only qualitatively estimated at low MAS rates. In principle, utilization of RNA with selectively labeled nitrogens in nucleobases and selective RNA deuteration could allow quantitative analysis of ^15^N *T*_1_ relaxation data for RNA at slow-to-moderate MAS rates. At the higher MAS rates (55 and 100 kHz), however, the effect of spin-diffusion on ^15^N *T*_1_ values becomes negligible, enabling initial characterization of site-specific RNA mobility via accurate measurement of ^15^N *T*_1_ time-constants. We observe a clear correlation between the measured ^15^N *T*_1_ relaxation time-constants and the different structural regions within the RNA molecule, which is readily interpretable in terms of the corresponding local flexibility of these regions. In particular, nitrogens in non-base-paired nucleotides in the k-turn and loop regions show significantly shorter relaxation time-constants, in agreement with the expected high mobility of these nucleotides, while those in base-paired nucleotides in the two stems and the k-turn are characterized by longer *T*_1_s, as expected for nucleotides with limited local flexibility.

The initial analysis of ^15^N *T*_1_ relaxation presented here demonstrates that ssNMR is a valuable technique for probing the dynamics and the conformational flexibility of RNA. We are currently working on further quantitative interpretation of ^15^N relaxation data and collection of ^13^C relaxation data at high MAS rate, which we anticipate will provide comprehensive picture of RNA dynamics.

## Acknowledgements

This work was supported by the Deutsche Forschungsgemeinschaft (DFG grants AM5157/3-1 and AM5157/6-1 to AM). We thank Prof. Teresa Carlomagno (University of Birmingham) for useful discussions.

## Author contributions

Conceptualization: A.M. Methodology: All authors. Investigation: All authors. Data curation: A.K.S. and A.M. Visualization: A.K.S. and A.M. Funding acquisition: A.M. Project administration: A.M. Supervision: A.M. Writing—original draft: A.K.S. and A.M. Writing—review and editing: All authors.

