## Supplementary Information for "Probing site-specific RNA dynamics by solid-state NMR spectroscopy"

**Abstract:** Knowledge of site-specific dynamics of biomolecules is necessary to understand their specific functions. Solid-state NMR is a uniquely powerful technique that can provide both structural and motional information about complex biomolecules. In recent years, nucleic acids have become increasingly studied by ssNMR, but as yet few RNA or DNA structures are available, with ssNMR-derived dynamics data on RNA rarer still. Here, we report the first systematic ssNMR study of $^{15}$N $T_1$ relaxation in RNA using straightforward nucleotide-type-specific and uniform labeling schemes. We observe clear correlation of the measured $^{15}$N $T_1$ relaxation time-constants with different structural elements in the RNA molecule, reflecting the distinct characteristics of their underlying motions. We anticipate that this novel approach can be further developed to provide a detailed and comprehensive picture of RNA dynamics in large biomolecular machines.

### Table of Contents

Figure S1. Schematic representations of NMR pulse-sequences and magnetization-transfer pathways. **2**
Table S1. Spectral acquisition parameters for each experiment at the three different MAS rates. **3**
Table S2. Cross-polarization parameters for each experiment at the three different MAS rates. **3**
Table S3. Processing parameters. **3**
Figure S2. $^{15}$N $T_1$ relaxation data at 16 kHz MAS and 600 MHz magnetic field. **5**
Figure S3. $^{15}$N $T_1$ relaxation data at 55 kHz MAS and 600 MHz magnetic field. **5**
Figure S4. $^{15}$N $T_1$ relaxation data at 100 kHz MAS and 850 MHz magnetic field. **6**
Figure S5. $^{15}$N $T_1$ relaxation data at 55 kHz MAS and 850 MHz magnetic field **6**
**Estimation of the impact of spin-diffusion on the extracted relaxation time-constants at 16 kHz MAS** **7**
Figure S6. Spin-diffusion in RNA nucleobases at 16 kHz MAS. **8**
Table S4. Spin diffusion exchange rates estimated from 10 s PDSD spectra. **9**
Table S5. Summary of all $T_1$ values for 26mer box C/D at different MAS rates **11**
Figure S7. Graphical representation of $T_1$ values measured for the 26mer box C/D RNA in complex with the L7Ae protein under varying MAS rates and magnetic field strengths11. **12**
**References** **12**

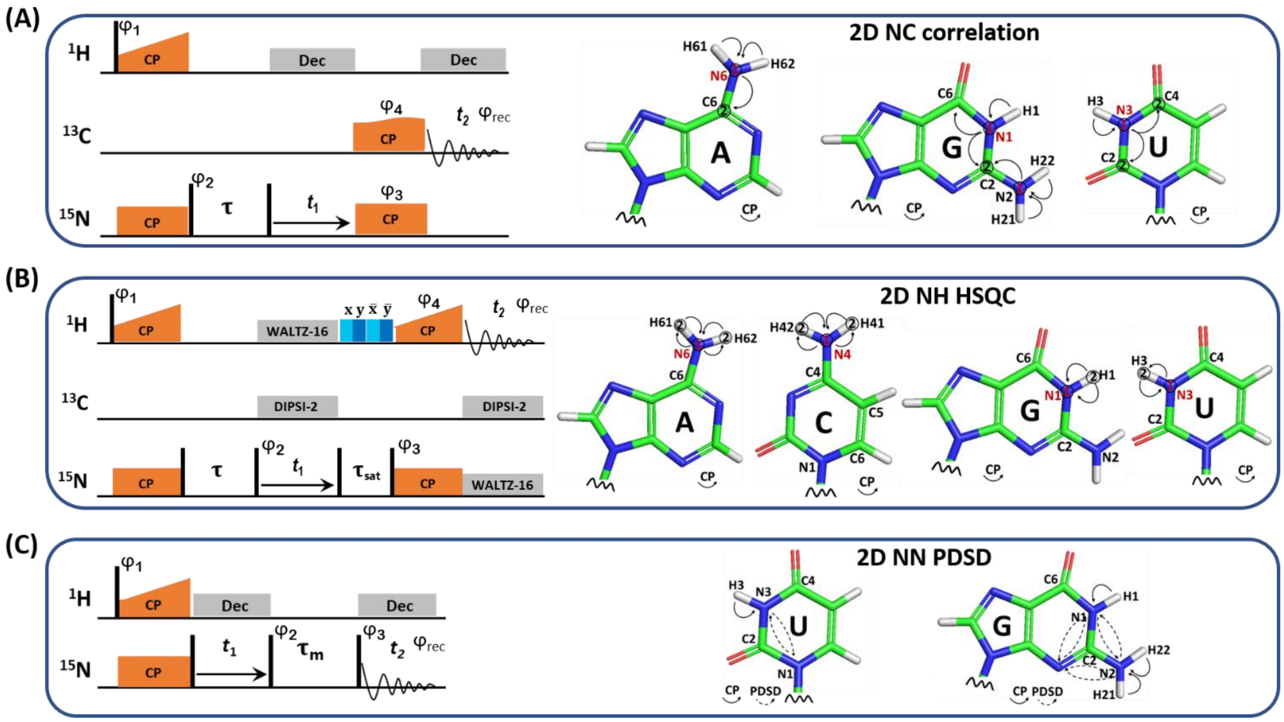

**Figure S1. Schematic representations of NMR pulse-sequences and magnetization-transfer pathways. (A)** 2D $^{13}$C,$^{15}$N correlation experiment for measurement of $^{15}$N $T_1$ relaxation time-constants at 16 kHz MAS. The initial $^{1}$H–$^{15}$N CP element is followed first by a ($\pi$/2–$\tau$–$\pi$/2) block comprising the relaxation delay $\tau$, and then an indirect $^{15}$N chemical-shift evolution period ($t_1$). Finally, magnetization is transferred by base-specific CP to $^{13}$C for detection ($t_2$). SPINAL-64[1] $^{1}$H decoupling (typically at a $B_1$ field-strength of 90 kHz) was applied during $t_1$ and $t_2$. **(B)** 2D $^{1}$H,$^{15}$N correlation experiment for measurement of $^{15}$N $T_1$ relaxation time-constants at 55 kHz and 100 kHz MAS. The initial $^{1}$H–$^{15}$N CP element is followed first by a ($\pi$/2–$\tau$–$\pi$/2) block comprising the relaxation delay $\tau$, and then an indirect $^{15}$N chemical-shift evolution period ($t_1$). Finally, after a water-saturation block, the magnetization is transferred by CP to $^{1}$H for detection ($t_2$). WALTZ-16 decoupling[2] was applied to $^{1}$H during $t_1$ (10 kHz $B_1$ field-strength) and to $^{15}$N during $t_2$ (10 kHz $B_1$ field-strength). DIPSI-2 decoupling[3] (20 kHz $B_1$ field-strength) was used for $^{13}$C decoupling during both $t_1$ and $t_2$. The MISSISSIPPI scheme[4] (40 kHz $B_1$ field-strength) was employed to suppress the residual $H_2O$ signal just prior to $^{1}$H detection. **(C)** 2D $^{15}$N,$^{15}$N PDSD experiment for measurement of spin-diffusion. The spectral acquisition parameters and precise parameters for the different cross-polarization steps are given in tables S1 and S2, respectively. The phase-programs were: **(A)** $\phi_1$ = y, –y; $\phi_2$ = 8(y), 8(–y); $\phi_3$ = 2(x), 2(–x); $\phi_4$ = 4(x), 4(–x); $\phi_{rec}$ = –x, x, x, –x, x, –x, –x, x, x, –x, –x, x, –x, x, x, –x; **(B)** $\phi_1$ = y, –y; $\phi_2$ = 8(x), 8(–x); $\phi_3$ = 2(x), 2(–x); $\phi_4$ = 4(y), 4(–y); $\phi_{rec}$ = y, –y, –y, y, 2(–y, y, y, –y), y, –y, –y, y; **(C)** $\phi_1$ = y, –y; $\phi_2$ = 8(x), 8(–x); $\phi_3$ = 2(x), 2(–x), 2(y), 2(–y); $\phi_{rec}$ = x, –x, –x, x, y, –y, –y, y, –x, x, x, –x, –y, y, y, –y.

**Table S1.** Spectral acquisition parameters for each experiment at the three different respective MAS rates and two magnetic fields.

| Experiment | Sample | Relaxation delay $\tau$, s | Acq. type | Spectral widths, kHz (ppm) | | $t_1$, ms | $t_2$, ms | Carrier freqs, ppm | PDSD mix time, s | Scans per point, ns | FID Size (F1, F2) | | Fn MODE |
|---|---|---|---|---|---|---|---|---|---|---|---|---|---|
| | | | | $^{15}$N | $^{13}$C | $^{15}$N | $^{13}$C | $^{15}$N, $^{13}$C | | | $^{15}$N | $^{13}$C | |
| **16 kHz MAS/600 MHz magnetic field** | | | | | | | | | | | | | |
| 2D NC N6-C6 | **A$^{lab}$ RNA** | 0, 3, 10, 20, 30, 50 | | 1.2 (20) | 45.5 (301) | 8.2 | 16.9 | 80, 100 | | 256 | 20 | 1536 | States-TPPI |
| 2D NC N1-C2 | **G$^{lab}$ RNA** | 0, 3, 5, 10, 20, 30, 40 | Inter-leaved | 3.0 (49) | 45.5 (301) | 7.9 | 16.9 | 90, 100 | | 128 | 48 | 1536 | States-TPPI |
| 2D NC N3-C4 | **U$^{lab}$ RNA** | 0, 3, 5, 10,15, 20, 30, 40, 50 | | 1.2 (20) | 45.5 (301) | 8.2 | 16.9 | 160, 100 | | 256 | 20 | 1536 | States-TPPI |

| | | | | | | | | | | | | | |
|---|---|---|---|---|---|---|---|---|---|---|---|---|---|
| 2D N1-N2 | **$G^{lab}$ RNA** | - | | 8.0 (131) | 24.5 (300) | 8 | 21 | 105, 105 | 1, 3, 5, 10, 20 | 128 | 128 | 1024 | States-TPPI |
| 2D N3-N1 | **$U^{lab}$ RNA** | - | | 4.0 (65) | 24.3 (400) | 8 | 21 | 160, 160 | 1, 3, 5, 10, 20 | 256 | 96 | 1024 | States-TPPI |
| **55 kHz MAS/600 MHz magnetic field** | | | | | | | | | | | | | |
| | | | | **$^{15}N$** | **1H** | **$^{15}N$** | **$^{1}H$** | **$^{15}N$, $^{1}H$** | | | **$^{15}N$** | **$^{1}H$** | |
| 2D NH N6-H61/62 | **$A^{lab}$ RNA** | 0, 5, 10, 15, 20, 30, 40 | Inter-leaved | 1.2 (20) | 6.1 (10) | 9.8 | 0.2 | 80, 6.7 | | 128 | 24 | 2048 | States-TPPI |
| 2D NH N1-H1 | **$G^{lab}$ RNA** | 0, 3, 5, 10, 20, 40, 60 | Non-inter-leaved | 2.4 (40) | 45.5 (75) | 9.8 | 22.5 | 79, 6.7 | | 128 | 48 | 2048 | States-TPPI |
| 2D NH N3-H3 | **$U^{lab}$ RNA** | 0, 5, 10, 15, 20, 30, 40, 60 | | 2.4 (40) | 45.5 (75) | 9.8 | 22.5 | 79, 6.1 | | 128 | 48 | 2048 | States-TPPI |
| **100 kHz MAS/850 MHz magnetic field** | | | | | | | | | | | | | |
| 2D NH A: N6-H61/62 C: N4-H41/42 G: N1-H1 U: N3-H3 | **Uniformly labeled RNA** | 0, 3, 5, 10, 20, 40, 60 | Inter-leaved | 10.3 (120) | 40.0 (47) | 4.8 | 25.6 | 117, 7.0 | | 32 | 100 | 2048 | States-TPPI |
| **55 kHz MAS/850 MHz magnetic field** | | | | | | | | | | | | | |
| 2D NH A: N6-H61/62 C: N4-H41/42 G: N1-H1 U: N3-H3 | **Uniformly labeled RNA** | 0, 3, 5, 10, 15, 20, 40, 60 | Inter-leaved | 8.5 (98.7) | 40.0 (47) | 5.9 | 25.6 | 117, 7.0 | | 32 | 100 | 2048 | States-TPPI |

**Table S2.** Cross-polarization parameters for each experiment at the three different MAS rates and two magnetic fields.

| **Experiment** | **Sample** | **CP transfer** | **CP contact time, ms** | **Type** | **Average RF strength, kHz** | | **RF pulse shape during CP** | |
|---|---|---|---|---|---|---|---|---|
| **16 kHz MAS/600 MHz magnetic field** | | | | | | | | |
| 2D NC N6-C6 | **$A^{lab}$ RNA** | **$^{1}H \rightarrow {}^{15}N$** | 0.6 | ZQ (n=1) | 56.3 | 40.7 | linear ramp up ±20% | rectangle |
| | | **$^{15}N \rightarrow {}^{13}C$** | 2.0 | ZQ (n=1) | 28.5 | 41.7 | tangential down ±10% | rectangle |
| 2D NC N1-C2 | **$G^{lab}$ RNA** | **$^{1}H \rightarrow {}^{15}N$** | 0.2 | ZQ (n=1) | 59.4 | 40.7 | linear ramp up ±20% | rectangle |
| | | **$^{15}N \rightarrow {}^{13}C$** | 3.0 | ZQ (n=1) | 28.5 | 42.5 | tangential down ±10% | rectangle |
| 2D NC N3-C4 | **$U^{lab}$ RNA** | **$^{1}H \rightarrow {}^{15}N$** | 0.6 | ZQ (n=1) | 50.2 | 35.1 | linear ramp up ±20% | rectangle |
| | | **$^{15}N \rightarrow {}^{13}C$** | 2.0 | ZQ (n=1) | 26.6 | 42.8 | tangential down ±10% | rectangle |
| 2D NN N1-N2 | **$G^{lab}$ RNA** | **$^{1}H \rightarrow {}^{15}N$** | 0.5 | ZQ (n=1) | 56.3 | 40.3 | linear ramp up ±20% | rectangle |
| 2D NN N3-N1 | **$U^{lab}$ RNA** | **$^{1}H \rightarrow {}^{15}N$** | 0.7 | ZQ (n=1) | 57.0 | 40.3 | linear ramp up ±20% | rectangle |
| **55 kHz MAS/600 MHz magnetic field** | | | | | | | | |
| 2D HN N6-H61/62 | **$A^{lab}$ RNA** | **$^{1}H \rightarrow {}^{15}N$** | 1.4 | ZQ (n=2) | 160.0 | 40.8 | linear ramp up ±30% | rectangle |
| | | **$^{15}N \rightarrow {}^{1}H$** | 1.2 | ZQ (n=2) | 40.8 | 151.2 | linear ramp up ±30% | rectangle |
| 2D HN N1-H1 | **$G^{lab}$ RNA** | **$^{1}H \rightarrow {}^{15}N$** | 1.6 | ZQ (n=2) | 158.6 | 40.9 | linear ramp up ±30% | rectangle |
| | | **$^{15}N \rightarrow {}^{1}H$** | 1 | ZQ (n=2) | 40.9 | 155.1 | linear ramp up ±30% | rectangle |
| 2D HN N3-H3 | **$U^{lab}$ RNA** | **$^{1}H \rightarrow {}^{15}N$** | 1.2 | ZQ (n=2) | 158.6 | 40.9 | linear ramp up ±30% | rectangle |
| | | **$^{15}N \rightarrow {}^{1}H$** | 1.2 | ZQ (n=2) | 40.9 | 162.4 | linear ramp up ±30% | rectangle |
| **100 kHz MAS/850 MHz magnetic field** | | | | | | | | |
| 2D NH A: N6-H61/62 C: N4-H41/42 G: N1-H1 U: N3-H3 | **Uniformly labeled RNA** | **$^{1}H \rightarrow {}^{15}N$** | 0.75 | DQ (n=1) | 78 | 25.2 | linear ramp up ±15% | rectangle |
| | | **$^{15}N \rightarrow {}^{1}H$** | 0.6 | ZQ (n=1) | 25.2 | 125.4 | linear ramp up ±15% | rectangle |
| **55 kHz MAS/850 MHz magnetic field** | | | | | | | | |
| 2D NH A: N6-H61/62 C: N4-H41/42 G: N1-H1 U: N3-H3 | **Uniformly labeled RNA** | **$^{1}H \rightarrow {}^{15}N$** | 0.75 | ZQ (n=1) | 99 | 41.5 | linear ramp up ±15% | rectangle |
| | | **$^{15}N \rightarrow {}^{1}H$** | 0.6 | ZQ (n=1) | 41.5 | 89 | linear ramp up ±15% | rectangle |

**Table S3**. Processing parameters.

| Experiment | Sample | Digital resolution | | Window function | | Box-width, ppm |
|---|---|---|---|---|---|---|
| | | $^{15}$N | $^{13}$C | $^{15}$N | $^{13}$C | $^{15}$N, $^{13}$C |
| **16 kHz MAS/600 MHz magnetic field** | | | | | | |
| 2D NC N6-C6 | **A$^{lab}$ RNA** | 128 | 8192 | GM –60Hz/0.2 | GM –60Hz/0.3 | 0.74, 0.78 |
| 2D NC N1-C2 | **G$^{lab}$ RNA** | 256 | 8192 | GM –60Hz/0.2 | GM –60Hz/0.3 | 0.97, 0.55 |
| 2D NC N3-C4 | **U$^{lab}$ RNA** | 128 | 8192 | EM 70Hz/0.2 | - | 0.98, 0.73 |
| 2D N1-N2 | **G$^{lab}$ RNA** | 1024 | 4096 | QSINE SSB=2 | QSINE SSB=2 | 0.49, 0.49 |
| 2D N3-N1 | **U$^{lab}$ RNA** | 1024 | 4096 | QSINE SSB=2 | QSINE SSB=2 | 0,65, 0.98 |
| | | $^{15}$N | $^{1}$H | $^{15}$N | $^{1}$H | $^{15}$N, $^{1}$H |
| **55 kHz MAS/600 MHz magnetic field** | | | | | | |
| 2D NH N6-H61/62 | **A$^{lab}$ RNA** | 128 | 8192 | GM –50Hz/0.1 | - | 1.56, 0.37 |
| 2D NH N1-H1 | **G$^{lab}$ RNA** | 256 | 8192 | EM 80Hz/0.1 | - | 1.1, 0.28 |
| 2D NH N3-H3 | **U$^{lab}$ RNA** | 256 | 8192 | EM 80Hz/0.1 | - | 1,26, 0.68 |
| **55 and 100 kHz MAS/850 MHz magnetic field** | | | | | | |
| 2D NH A: N6-H61/62 C: N4-H41/42 G: N1-H1 U: N3-H3 | **Uniformly labeled RNA** | 256 | 16384 | GM –60Hz/0.3 | QSINE SSB=3 | 1.5, 0.2 |

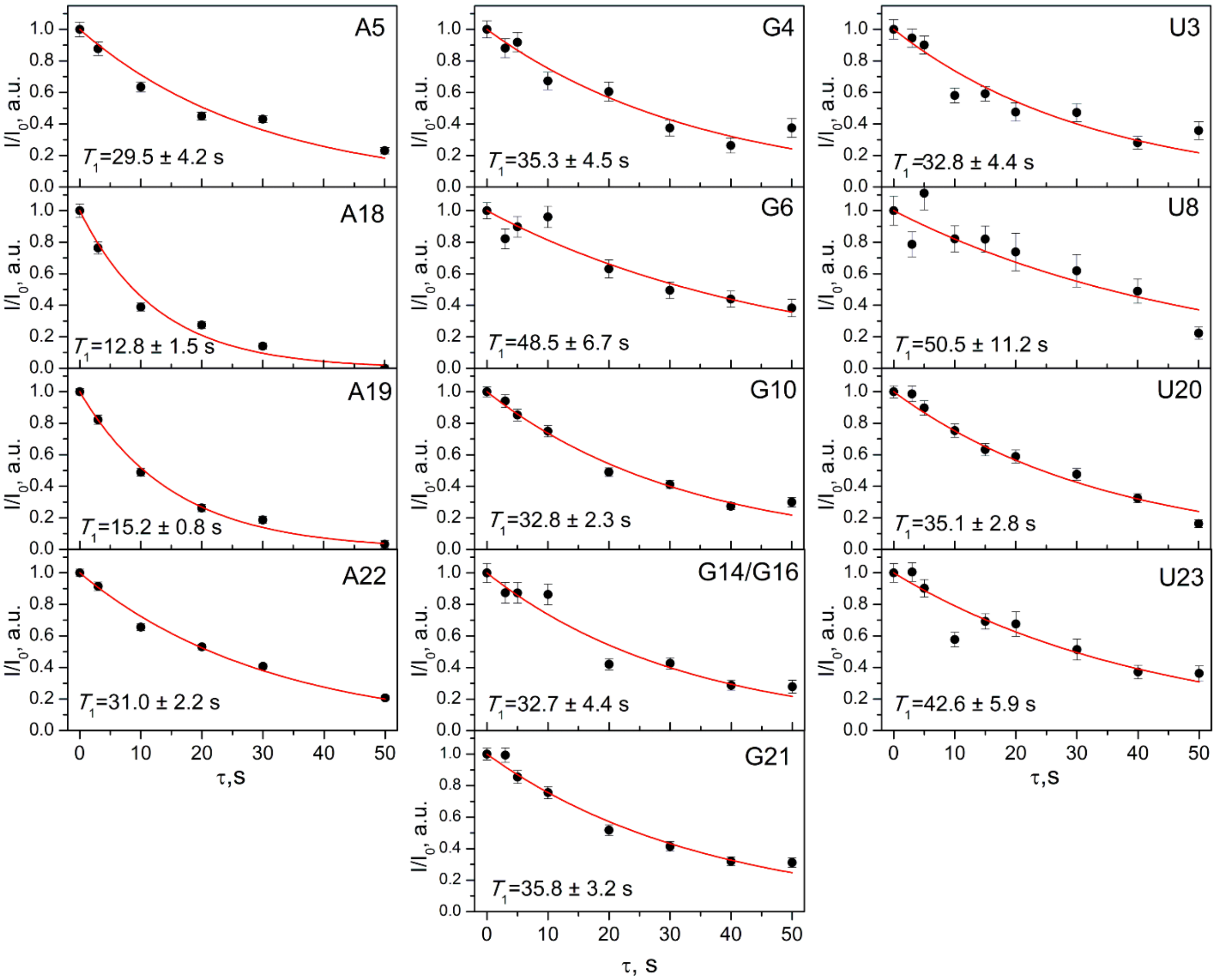

**Figure S2. $^{15}$N $T_1$ relaxation data at 16 kHz MAS and 600 MHz magnetic field.** Experimental relaxation curves for all analyzed nucleotides at 16 kHz MAS and 600 MHz field-strength. Mono-exponential decay fits are shown in red.

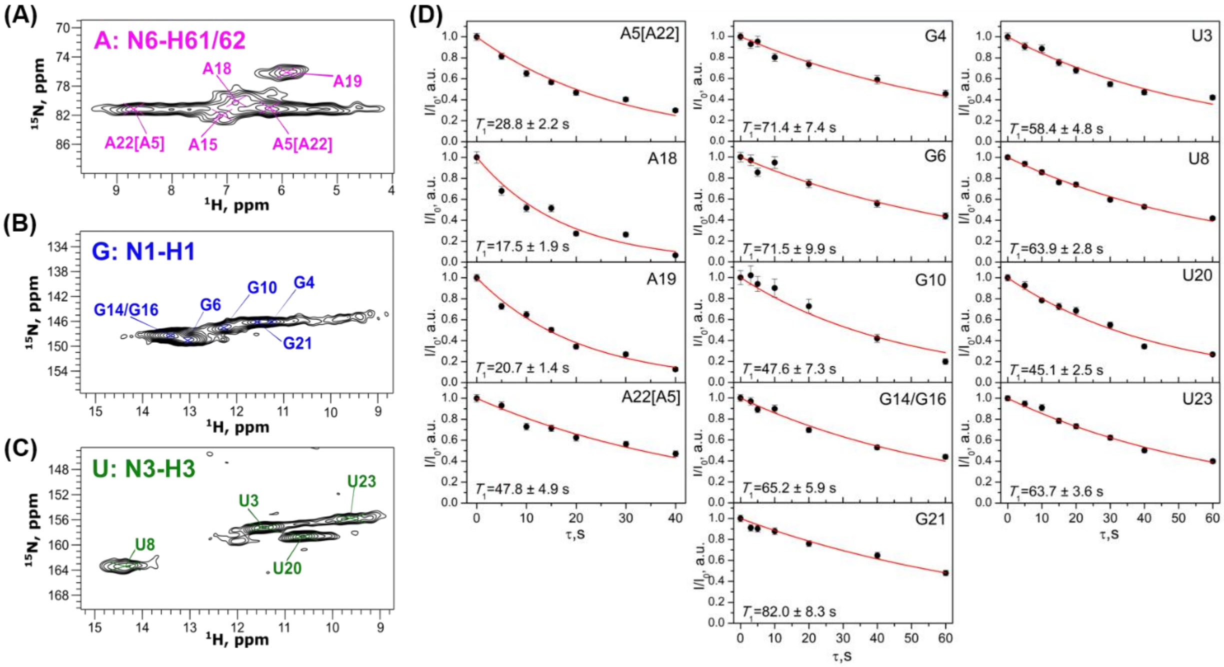

**Figure S3. $^{15}$N $T_1$ relaxation data at 55 kHz MAS and 600 MHz magnetic field.** 2D $^{1}$H,$^{15}$N spectra of adenosines **(A)**, guanosines **(B)** and uridines **(C)** of 26mer box C/D RNA in complex with L7Ae protein. **(D)** Experimental relaxation curves for all analyzed nucleotides. Mono-exponential decay fits are shown in red. All data was acquired at 55 kHz MAS and a field-strength of 600 MHz.

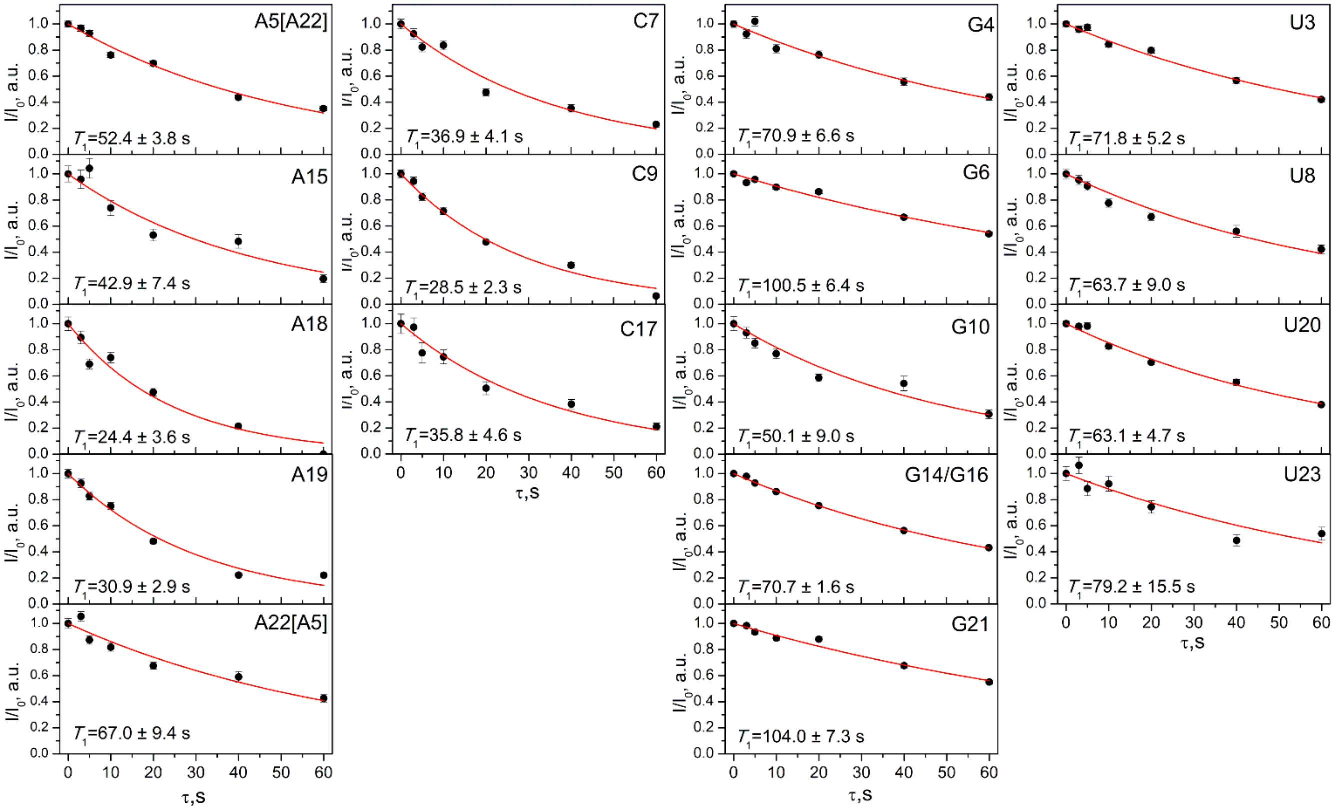

**Figure S4. $^{15}$N $T_1$ relaxation data at 100 kHz MAS and 850 MHz magnetic field.** Experimental relaxation curves for all analyzed nucleotides. Mono-exponential decay fits are shown in red. All data was acquired at 100 kHz MAS and a field-strength of 850 MHz.

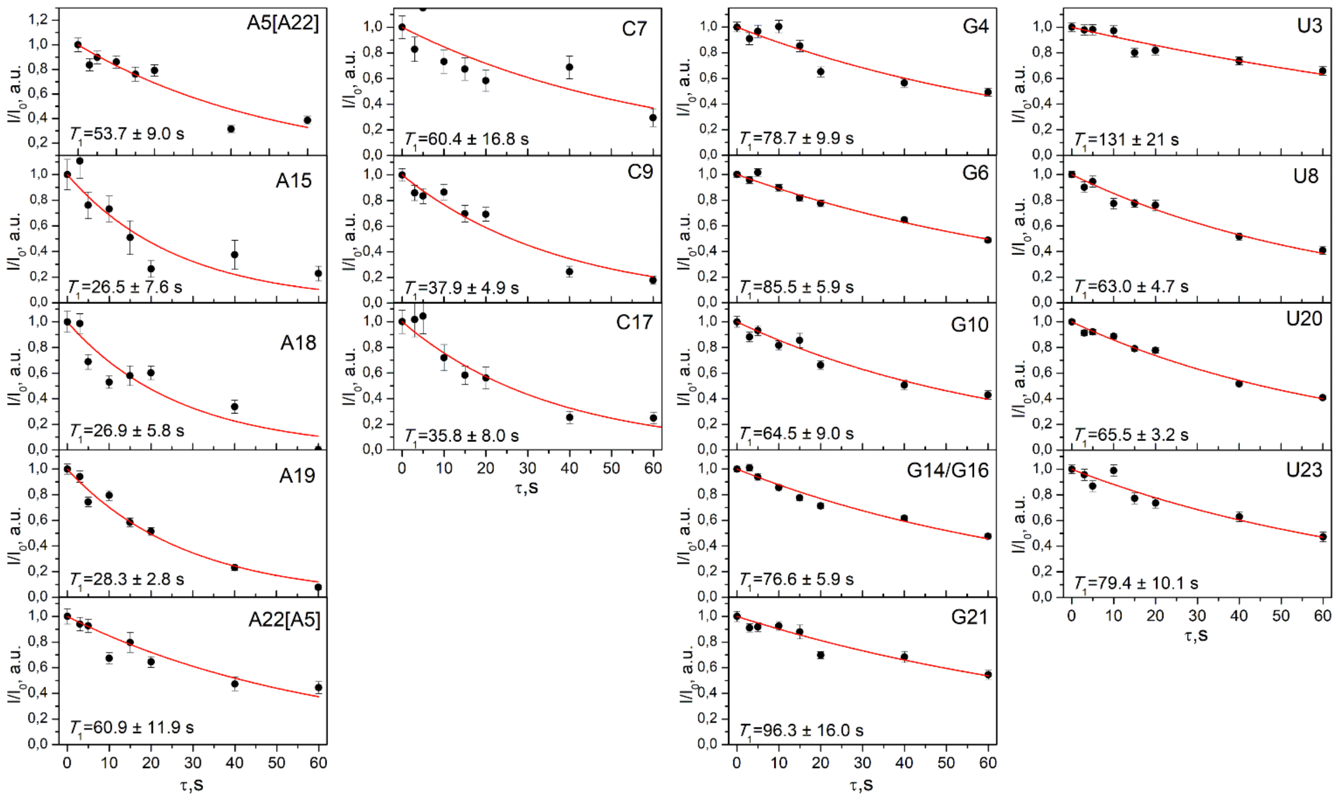

**Figure S5. $^{15}$N $T_1$ relaxation data at 55 kHz MAS and 850 MHz magnetic field.** Experimental relaxation curves for all analyzed nucleotides. Mono-exponential decay fits are shown in red. All data was acquired at 100 kHz MAS and a field-strength of 850 MHz.

#### Estimation of the impact of spin-diffusion on the extracted relaxation time-constants at 16 kHz MAS

The impact of spin-diffusion on the measured relaxation time-constants at 16 kHz was estimated according to the procedure of Giraud et al.[5] The time-evolution of the longitudinal magnetization of a two-spin system (for example, nitrogens N1 and N2 in guanosine), considering both auto-relaxation and magnetization exchange due to PDSD, is described by the following pair of coupled differential equations:[5]

$$\frac{\partial N1}{\partial t} = -R_1^{N1}(N1(t) - M_0) - \sigma(N1(t) - N2(t)),$$
$$\frac{\partial N2}{\partial t} = -R_1^{N2}(N2(t) - M_0) - \sigma(N2(t) - N1(t)), \qquad (2)$$

where *N1*(*t*) and *N2*(*t*) are the time-dependent magnetizations of nitrogens N1 and N2 respectively, with true auto-relaxation rates $R_1^{N1} = 1/T_1^{N1}$ and $R_1^{N2} = 1/T_1^{N2}$, respectively. The forward and backward exchange rates are assumed to be equal, i.e. $\sigma_{12} = \sigma_{21} = \sigma$. $M_0$ represents the nitrogen magnetization at thermal equilibrium; this is assumed to be the same for both nitrogens. Using an appropriate change of variables, this pair of equations set can be solved analytically, and the resulting theoretical relaxation profiles $N_{\text{theor}}(t)$ for nitrogens N1 and N2 can be written as:[5-6]

$$N1_{\text{theor}}(t) = M_0 + \frac{1}{2\sqrt{\left((R_1^{N1}+\sigma)-(R_1^{N2}+\sigma)\right)^2+4\sigma^2}} e^{-\frac{1}{2}t\left(\sqrt{\left((R_1^{N1}+\sigma)-(R_1^{N2}+\sigma)\right)^2+4\sigma^2}+(R_1^{N1}+\sigma)+(R_1^{N2}+\sigma)\right)} \times$$
$$\left(e^{t\sqrt{\left((R_1^{N1}+\sigma)-(R_1^{N2}+\sigma)\right)^2+4\sigma^2}} \left(x_0\left(-(R_1^{N1}+\sigma)\right) + x_0(R_1^{N2}+\sigma) + 2y_0\sigma\right) + \right.$$
$$x_0\sqrt{\left((R_1^{N1}+\sigma)-(R_1^{N2}+\sigma)\right)^2+4\sigma^2}\, e^{t\sqrt{\left((R_1^{N1}+\sigma)-(R_1^{N2}+\sigma)\right)^2+4\sigma^2}} + x_0\sqrt{\left((R_1^{N1}+\sigma)-(R_1^{N2}+\sigma)\right)^2+4\sigma^2} + x_0(R_1^{N1}+\sigma) -$$
$$\left. x_0(R_1^{N2}+\sigma) - 2y_0\sigma\right),$$

$$N2_{\text{theor}}(t) = M_0 + \frac{1}{2\sqrt{\left((R_1^{N1}+\sigma)-(R_1^{N2}+\sigma)\right)^2+4\sigma^2}} e^{-\frac{1}{2}t\left((R_1^{N1}+\sigma)+(R_1^{N2}+\sigma)+\sqrt{\left((R_1^{N1}+\sigma)-(R_1^{N2}+\sigma)\right)^2+4\sigma^2}\right)} \times \left(-(R_1^{N1}+\sigma)y_0 + (R_1^{N2}+\sigma)y_0 - \right.$$
$$2x_0\sigma + e^{t\sqrt{\left((R_1^{N1}+\sigma)-(R_1^{N2}+\sigma)\right)^2+4\sigma^2}}\left((R_1^{N1}+\sigma)y_0 - (R_1^{N2}+\sigma)y_0 + 2x_0\sigma\right) + y_0\sqrt{\left((R_1^{N1}+\sigma)-(R_1^{N2}+\sigma)\right)^2+4\sigma^2} +$$
$$\left. e^{t\sqrt{\left((R_1^{N1}+\sigma)-(R_1^{N2}+\sigma)\right)^2+4\sigma^2}} y_0\sqrt{\left((R_1^{N1}+\sigma)-(R_1^{N2}+\sigma)\right)^2+4\sigma^2}\right), \qquad (3)$$

where $x_0 = N1(0) - M_0$ and $y_0 = N2(0) - M_0$ represent the deviations of the magnetizations of N1 and N2 from their equilibrium values at time-zero.

Assuming that the $T_1$ values measured at 55 kHz MAS represent good estimates of the true auto-relaxation $T_1$ values (i.e. assuming that spin-diffusion is negligible at 55 kHz MAS), it is possible to use the equations above to simulate expected intensity-decay profiles at 16 kHz MAS; the simulation parameters comprise the experimental $T_1$ values measured at 55 kHz MAS, estimates of the initial magnetizations at 16 kHz MAS and the experimentally measurable exchange rate-constants measured at 16 kHz MAS, together with a range of possible values for experimentally inaccessible parameters. The resulting simulated intensity-decay profiles, which incorporate the effects of both relaxation and non-negligible spin-diffusion, can be fitted to the mono-exponential decay-function to yield apparent $T_1$ values at 16 kHz MAS, $T_1^{\text{app,sim}}$. The deviation between these apparent $T_1$ values obtained by fitting the simulated profiles and the true $T_1$ values used as input parameters for the simulation provides an indication of the degree of error introduced by attempting to fit the experimental data at 16 kHz MAS to a mono-exponential decay-function; if the implemented model is appropriate, these $T_1^{\text{app,sim}}$ values should be in good agreement with the corresponding apparent $T_1$ values obtained by fitting the experimental data at 16 kHz MAS ($T_1^{\text{app,exp}}$).

Measuring exchange rates due to spin diffusion in RNA is challenging since the nitrogen-density in the RNA nucleobases is higher than in proteins but many of these nitrogens are not protonated. The set of nitrogen-pairs in nucleobases for which magnetization-exchange by spin-diffusion may be problematic in the context of

$^{15}$N $T_1$ measurements is comprised by those nitrogen-pairs whose separation is less than an estimated spin-diffusion cut-off of 3.0 Å and in which one or both of the nitrogen atoms is protonated.

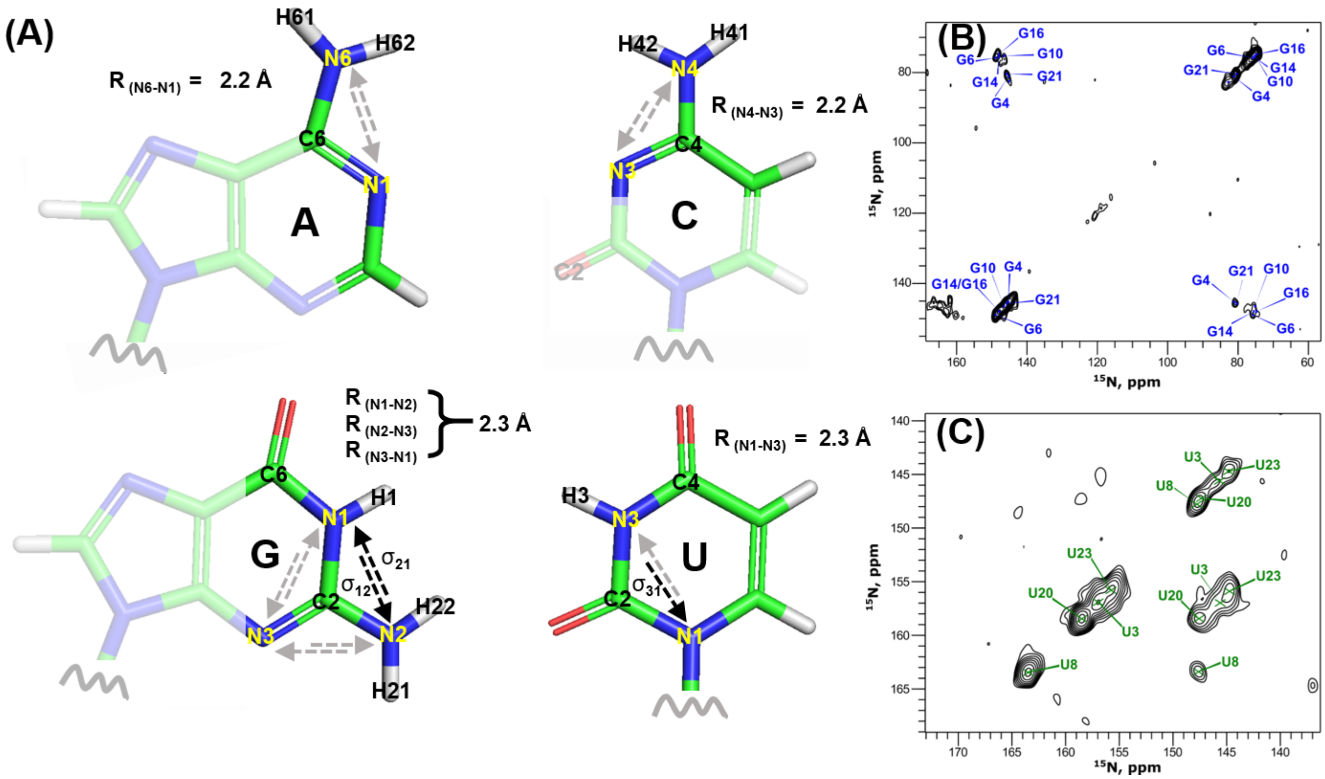

**Figure S6. Spin-diffusion in RNA nucleobases at 16 kHz MAS. (A)** The structures of the four RNA nucleobases; dashed arrows illustrate magnetization exchange between nitrogens within a 3.0 Å cut-off. Measured and not-measured exchange rates are shown as black and gray arrows, respectively. 2D $^{15}$N,$^{15}$N PDSD spectra of **(B)** G$^{lab}$-26mer RNA and **(C)** U$^{lab}$-26mer RNA at 10 s PDSD mixing-time. All data was acquired on nucleotide-type-specifically $^{13}$C,$^{15}$N-labeled 26mer box C/D RNA in complex with unlabeled L7Ae protein at a field-strength of 600 MHz.

An extra complication is that, in addition to intra-nucleotide N–N spin-diffusion, inter-strand spin-diffusion between base-paired nucleotides may also result in non-negligible magnetization exchange between the proximal nitrogen atoms. For example, the N1(G)–N3(C) and N3(U)–N1(A) distances across the Watson-Crick G:C and A:U base-pairs are 2.9 Å and 3.1 Å, respectively.[7] All of these considerations mean that the quantitative assessment of the impact of spin-diffusion on the measurement of $^{15}$N $T_1$s in RNA is extremely challenging. Nevertheless, we have derived ranges for apparent $^{15}$N $T_1$ relaxation time-constants at 16 kHz MAS ($T_1^{app,sim}$) in adenosines, guanosines and uridines at 16 kHz MAS according to the procedure described above. All calculations have been performed in Matlab.

**Table S4.** Spin diffusion exchange rates estimated from 10 s PDSD spectra.

| Residue | N1–N2 exchange rate, s$^{-1}$ | | | | Residue | N3–N1 exchange rate, s$^{-1}$ |
|---|---|---|---|---|---|---|
| | $\sigma_{12}$ | $\sigma_{21}$ | $\sigma$ | | | $\sigma_{31}$ |
| G4 | 0.017 | 0.025 | 0.021 | | U3 | 0.027 |
| G6 | 0.015 | 0.016 | 0.016 | | U8 | 0.015 |
| G10 | 0.007 | 0.009 | 0.008 | | U20 | 0.022 |

| | | | | | | |
|---|---|---|---|---|---|---|
| G14 | 0.011 | 0.009 | 0.010 | | U23 | 0.053 |
| G16 | 0.008 | 0.014 | 0.011 | | | |
| G21 | 0.009 | 0.031 | 0.019 | | U3, U8, U20 | 0.021 |
| All Gs | 0.011 | 0.017 | 0.014 | | All Us | 0.029 |

*Adenosines*

In adenosines, nitrogens N1 and N6 are located within the 3.0 Å cut-off, being separated by 2.2 Å (Fig. S6A). Based on the appearance of 1D $^{15}N$ CP spectra, the initial magnetization on nitrogen N1 was estimated to be 20% of that on nitrogen N6; additional calculations with values of 10–40% have been performed, which lead to a wider range of estimated $T_1^{app}$. Exchange-rates for adenosines were not measurable due to the severe spectral overlap but were assumed to be similar to those of uridines (0.015 $s^{-1}$); calculations were performed over the range 0.005–0.040 $s^{-1}$. Finally, it was assumed that $T_1$(N6) < $T_1$(N1) and calculations were performed with $T_1$(N1) in the range of 150–250% $T_1$(N6).

*Guanosines*

The distances between protonated nitrogens N1 and N2 and non-protonated nitrogen N3 in guanosines are 2.3 Å, which means that it is necessary to solve the corresponding set of three coupled differential equations similar to those of equation-set (2). The initial magnetization on nitrogen N3 was estimated to be 10% of that on nitrogen N1, while initial magnetization on N2 was estimated to be 125% of that on nitrogen N1. Additional simulations were performed over the range of N2 magnetization of 100–170% N1. Exchange-rates $\sigma_{12}$ and $\sigma_{21}$ were calculated from 2D $^{15}N$,$^{15}N$ PDSD spectra acquired at 10 s mixing-time as $\sigma = \frac{I_{\text{cross-peak}}/I_{\text{diagonal}}}{\text{PDSD mixing time}}$ and were found to be different in several instances (Table S4), so that average values (calculated as $\sigma = (\sigma_{12} + \sigma_{21})/2$) have been used in the calculations. Due to substantial overlap of resonances in 2D $^{15}N$,$^{15}N$ PDSD spectra (Figure S6B), the measured exchange-rates are prone to errors, so the calculations were performed over the range 0.005–0.040 $s^{-1}$. The exchange-rates $\sigma_{13}/\sigma_{31}$ and $\sigma_{23}/\sigma_{32}$ were not measurable due to spectral overlap and calculations with values in the range 0.010–0.020 $s^{-1}$ have been performed, assuming slower exchange rates than for $\sigma_{12}/\sigma_{21}$ due to the absence of protons attached to nitrogen N3. Also, it was assumed that $T_1$(N3) > $T_1$(N1) > $T_1$(N2), and calculations with $T_1$(N2) in the range of 50–80% $T_1$(N1) and with two different values for $T_1$(N3) (120 & 150 s) were performed.

*Uridines*

In uridines, the distance between nitrogens N1 and N3 is 2.3 Å (Figure S6A). The initial magnetization on nitrogen N1 was estimated to be 35% of that on nitrogen N3; additional calculations with values of 10–40% N3 have been performed. The exchange-rates $\sigma_{31}$ were calculated from 2D $^{15}N$,$^{15}N$ PDSD spectra acquired with 10 s mixing-time (Figure S5C), additional calculations were performed over the range 0.005–0.040 $s^{-1}$. Also, it was assumed that $T_1$(N1) > $T_1$(N3) and calculations were performed with $T_1$(N1) in the range of 125–200% $T_1$(N3).

The simulation/fitting procedure provided the ranges of $T_1^{app,sim}$ values at 16 kHz MAS calculated on the basis of experimental $T_1$ values measured at 55 kHz MAS, which are given in table S5 and visualized in Figure S7A. The overall picture that emerges is that apparent relaxation time-constants for slowly relaxing imino nitrogens at 16 kHz MAS — as obtained by fitting intensity-decay profiles to a mono-exponential decay-function — can be up to 30% shorter than the true $T_1$ values (as measured at 55 kHz MAS). Moreover, most of these simulated apparent $T_1$ values are in very good agreement with experimental apparent $T_1$ values at 16 kHz MAS. We could not obtain good agreement for some of the data-sets (e.g. U3, G21, A22). Several factors are likely to contribute to the discrepancies in these cases: (i) the simplified experimental model; (ii) lack of experimental data and (iii) unknown systematic errors in the acquired experimental data, as might arise due to field-drift and/or temperature instabilities over the long experimental duration.

**Table S5.** Summary of all $T_1$ values for 26mer box C/D RNA in complex with L7Ae protein at different MAS rates and different magnetic fields. * indicates that the nucleotides peaks are partially overlapped in the NC correlation spectra at 16 kHz MAS (600 MHz), as illustrated in Figure 1D; # denotes that the A5/A22 peaks were completely overlapped by each other, as shown in Figures 2 and S3A-C.

| Region | Residue (N-atom) | $T_1^{app,exp}$ 16 kHz MAS (600 MHz) | | Residue (N-atom) | $T_1^{app,sim}$ 16 kHz MAS (600 MHz) | $T_1^{exp}$ 55 kHz MAS (600 MHz) | $T_1^{exp}$ 100 kHz MAS (850 MHz) | $T_1^{exp}$ 55 kHz MAS (850 MHz) |
|---|---|---|---|---|---|---|---|---|
| | G1 | | | G1 | | - | - | |
| Non-canonical stem | C2 | | | C2 | | - | - | |
| | U3 (N3) | 32.8 ± 4.4 | | U3 | 39–54 | 58.4 ± 4.8 | 71.8 ± 5.2 | 131 ± 21 |
| Kink turn | G4* (N1) | 35.3 ±4.5 | | G4 | 34–62 | 71.4 ± 7.4 | 70.9 ± 6.6 | 78.7 ± 9.9 |
| | A5 (N6) | 29.5 ± 4.2 | | A5[A22] (N6) | 22–29 | 28.2 ± 2.2 | 52.4 ± 3.8 | 53.7 ± 9.0 |
| Canonical stem | G6 (N1) | 48.5 ± 6.7 | | G6 | 35–63 | 71.5 ± 9.9 | 100.5 ± 6.4 | 85.5 ± 5.9 |
| | C7 (N4) | - | | C7 | | - | 36.9 ± 4.1 | 60.4 ± 16.8 |
| | U8 (N3) | 50.5 ± 1.2 | | U8 | 42–59 | 63.9 ± 2.8 | 63.7 ± 9.0 | 63.0 ± 4.7 |
| | C9 | - | | C9 | | - | 28.5 ± 2.3 | 37.9 ± 4.9 |
| Loop | G10* (N1) | 32.8 ± 2.3 | | G10 | 24–44 | 47.6 ± 7.3 | 50.1 ± 9.0 | 64.5 ± 9.0 |
| | A11 | - | | A11 | | - | - | - |
| | A12 | - | | A12 | | - | - | - |
| | A13 | - | | A13 | | - | - | - |
| Canonical stem | G14/G16 (N1) | 32.7 ± 4.4 | | G14/G16 (N1) | 31–58 | 65.2 ± 5.9 | 70.7 ± 1.6 | 76.6 ± 5.9 |
| | A15 | - | | A15 | | - | 42.9 ± 7.4 | 26.5 ± 7.6 |
| | G16/G14 (N1) | 32.7 ± 4.4 | | G16/G14 (N1) | 31–58 | 65.2 ± 5.9 | 70.7 ± 1.6 | 76.6 ± 5.9 |
| | C17 | - | | C17 (N4) | | - | 35.8 ± 4.6 | 35.8 ± 8.0 |
| Kink turn | A18 (N6) | 12.8 ± 1.5 | | A18 | 14–18 | 17.5 ± 1.9 | 24.4 ± 3.6 | 26.9 ± 5.8 |
| | A19 (N6) | 15.2 ± 0.8 | | A19 | 16–22 | 20.7 ± 1.4 | 30.9 ± 2.9 | 28.3 ± 2.8 |
| | U20 (N3) | 35.1± 2.8 | | U20 | 32–43 | 45.1 ± 2.5 | 63.1 ± 4.7 | 65.5 ± 3.2 |
| | G21* (N1) | 35.8 ± 3.2 | | G21 | 39–71 | 82.0 ± 8.3 | 104.0 ± 7.3 | 96.3 ± 16 |
| | A22 (N6) | 31.0 ± 2.2 | | A22[A5] | 34–46 | 47.8± 4.9 | 67.0 ± 9.4 | 60.9 ± 11.9 |
| Non-canonical stem | U23 (N3) | 42.6 ± 5.9 | | U23 | 42–59 | 63.7 ± 3.6 | 79.2 ± 15.5 | 79.4 ± 10.1 |
| | G24 | - | | G24 | | - | - | |
| | U25 | - | | U25 | | - | - | |

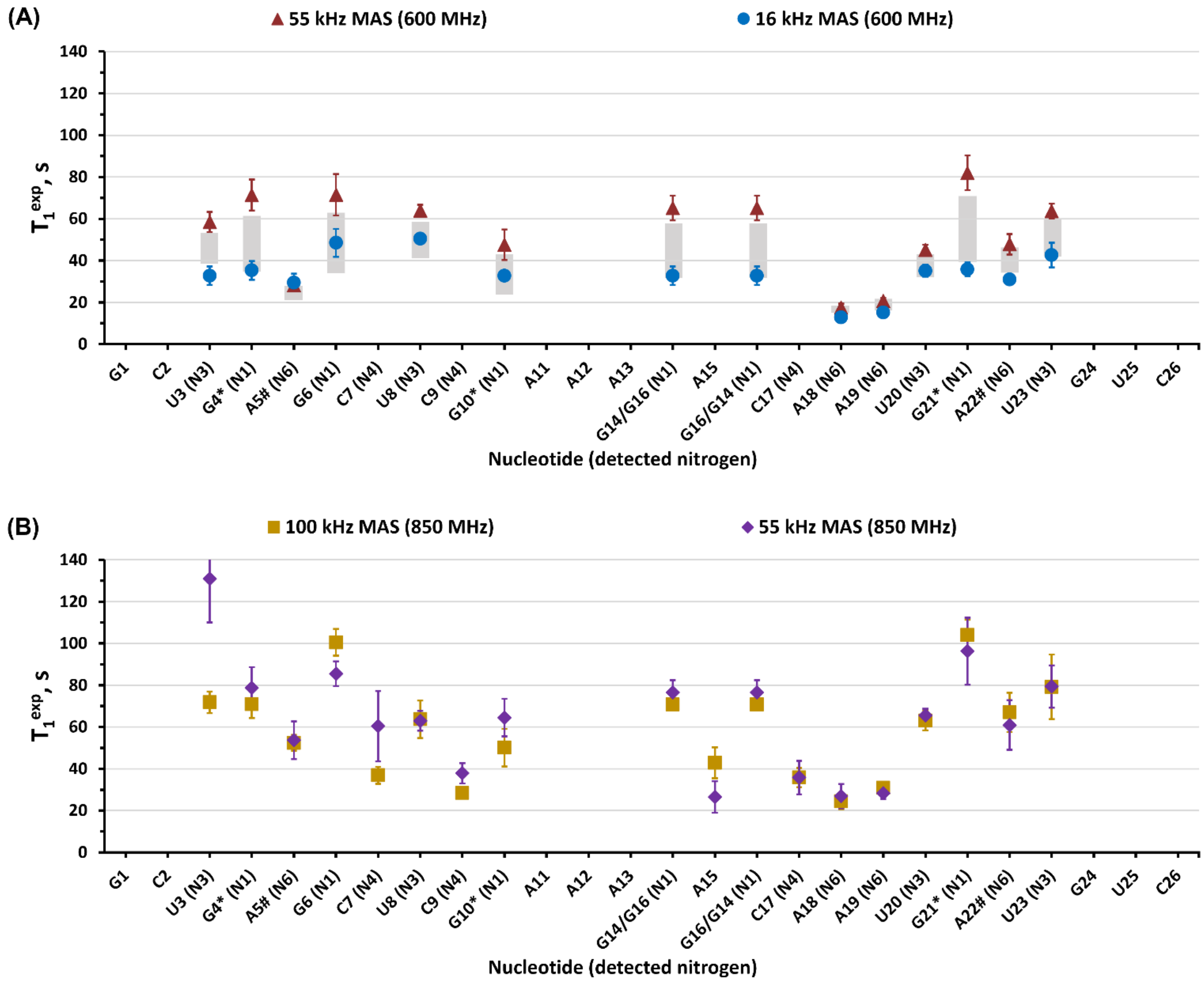

**Figure S7. Graphical representation of $T_1$ values measured for the 26mer box C/D RNA in complex with the L7Ae protein under varying MAS rates and magnetic field strengths. (A)** Comparison of $T_1$ values obtained at 16 kHz MAS ($T_1^{app,exp}$, blue circles) and 55 kHz MAS (red triangles) at 600 MHz. The ranges of theoretically estimated $T_1$ values ($T_1^{app,sim}$) are depicted as grey bars. **(B)** Comparison of experimental $T_1$ values at 55 kHz MAS (blue iamonds) and 100 kHz MAS (orange squares) at 850 MHz. (*) indicate nucleotides with partially overlapping peaks in the NC correlation spectra at 16 kHz MAS (600 MHz), as shown in Figure 1D. (#) denote complete overlap of the A5/A22 peaks, as illustrated in Figures S3(A–C) and 2.
